# Acoustic remote sensing with deep learning enables non-invasive estimation of seabird nest density

**DOI:** 10.64898/2026.01.13.699273

**Authors:** Francesca Terranova, Lorenzo Todaro, Xavier Forte, Katrin Ludynia, Deon Geldenhuys, Nicolas Mathevon, David Reby, Livio Favaro

## Abstract

Passive Acoustic Monitoring (PAM) has advanced ecological research by enabling non-invasive recordings of wildlife vocalizations that provide insight into species presence, behavior, and reproductive activity. This remote-sensing approach is particularly valuable for species breeding in concealed habitats or remote areas where visual surveys are challenging. The Critically Endangered African Penguin, burrow-nesting seabird, exemplifies this challenge. Its highly vocal breeding behavior makes it an ideal case study for evaluating passive acoustic monitoring as a low-disturbance approach to estimating nest density.

To evaluate this, we deployed Autonomous Recording Units at multiple sampling points across the Stony Point Penguin Colony, capturing soundscapes spanning different nest densities and environmental conditions. We then developed an automated detector for Ecstatic Display Songs (EDS), the species’ characteristic territorial song, using a Convolutional Neural Network trained on a multi-source dataset covering several breeding seasons, diverse acoustic environments, and both *in situ* and *ex situ* recordings. The model achieved high recall and precision and remained robust across diverse environmental conditions, supporting the use of heterogeneous training datasets for reliable bioacoustics detection.

Using the automated EDS detections, we then investigated how vocal activity peaks relate to local nest density. A Generalized Additive Model revealed that EDS peaks strongly predicted nest density, with a nonlinear increase that plateaued at high calling rates. Importantly, models trained in one breeding season generalized well to the next.

In conclusion, by integrating PAM with deep learning, this study provides a scalable, low-disturbance framework for estimating penguin nest density from soundscape data, supporting colony rangers in monitoring penguin colonies.

**Highlights:**

- Large-scale acoustic monitoring captured the soundscape of a critically endangered seabird species.
- Deep-learning-based acoustic detection reliably identified key breeding vocalizations in field recordings.
- Peak vocal activity strongly and nonlinearly predicted active nest density across sampling points.
- The vocal-nest relationship increases rapidly and plateaued at high levels of vocal activity.
- This approach enables scalable, low-disturbance monitoring of seabird nest density.

## 1. Introduction

Biodiversity loss is among the most pressing global challenges, accelerated by human activities that threaten ecosystems worldwide (Díaz et al., 2006, 2019; Johnson et al., 2017; Pereira et al., 2012; Werdelin & Lewis, 2013). This urgency has intensified the need for non-invasive, scalable monitoring tools that provide continuous ecological information while minimizing disturbance. Advances in satellite imagery, camera traps, and acoustic sensors have collectively enabled a new generation of remote sensing approaches for biodiversity monitoring (Browning et al., 2017; Gibb et al., 2019; McGeady et al., 2023).

To address limitations inherent to optical and aerial platforms, particularly their reduced effectiveness in visually obstructed habitats, ground-based acoustic remote sensing using *in situ* devices has become more widely used. Passive Acoustic Monitoring exemplifies this shift: it uses Autonomous Recording Units (ARUs) to capture environmental soundscapes at high temporal resolution (Ross et al., 2023; Sugai et al., 2019). Because sound propagates through vegetation and around obstacles, acoustic remote sensing can reveal ecological processes, including presence, behavior and reproductive activity, that remain invisible to optical or thermal sensors (Boullhesen et al., 2021; Browning et al., 2017; Gibb et al., 2019). PAM therefore provides a powerful, cost-effective complement to traditional remote-sensing platforms, offering continuous, minimally intrusive monitoring across challenging environments.

The advent of battery-powered ARUs in the late 1990s enabled long-term, non-invasive wildlife monitoring, proving effective across applications ranging from rare species detection to habitat-level biodiversity assessment (Blumstein et al., 2011; Gibb et al., 2019; Sugai et al., 2019). Recent advances in data storage, programmable recording schedules, and battery technology nowadays make it feasible to deploy ARUs across broad spatial scales. Parallel advances in analytical methods have transformed how animal vocalizations can be detected and analyzed.

Rapid progress in Artificial Intelligence has transformed the analysis of acoustic data. Deep learning, and particularly Convolutional Neural Networks (CNNs), has become central to automated species detection, excelling at extracting complex frequency patterns from spectrograms and reducing the need for manual annotation (LeCun et al., 2015; Sharma et al., 2023). Reliable detections are a prerequisite for ecological inference, once vocalizations are identified in continuous recordings, they can be summarized into vocal activity indices that describe vocal activity.

Using acoustics to estimate animal density requires methods that accommodate species-specific ecology and behavioral variability. Classical frameworks such as cue counting assume that vocalization rates scale predictably with individual numbers (Marques et al., 2013; Pérez-Granados & Traba, 2021). However, these assumptions can break down in species whose calling behavior is influenced by time of day, social interactions, or environmental conditions (Hutschenreiter et al., 2024; Navine et al., 2024; Upham-Mills et al., 2020). To address these challenges, recent work has developed more flexible vocal activity indices, including call density (Navine et al., 2024), Vocal Activity Rate, Detection Rate, and Maximum Count per Minute (Hutschenreiter et al., 2024). These approaches highlight the need for species-specific calibration when translating vocal output into meaningful ecological information.

Seabirds represent an ideal taxon where PAM might assist with population monitoring, thereby providing information essential for conservation management. They are among the most threatened vertebrate groups, with nearly half of all species at risk of extinction (Díaz et al., 2019), and many populations inhabit remote, inaccessible areas, often nesting in visually obstructed habitats. Penguins exemplify both the monitoring challenges and the inherent vulnerability of seabird species: 11 of the 18 recognized species (Gimeno et al., 2024; Hickcox et al., 2019) are threatened, with widespread declines driven by overfishing, climate change, invasive species, and habitat disturbance (Boersma et al., 2020; Phillips et al., 2016; Rodríguez et al., 2019; Trathan et al., 2015; Young & Ballance, 2023). Among them, the African Penguin (*Spheniscus demersus*) has undergone a dramatic population decline since the 1950s (Sherley et al., 2024) and is now classified as Critically Endangered by the International Union for Conservation of Nature (IUCN; BirdLife International, 2024, *Spheniscus demersus*). The decline has been driven by multiple pressures, including climate change–induced habitat degradation (Crawford et al., 2000), heat waves that reduce breeding success (Welman et al., 2024), and human impacts such as competition with commercial fisheries and coastal habitat disturbance (Crawford et al., 2019; Gimeno et al., 2024; Sherley et al., 2024). Monitoring African Penguin colonies is challenging: many colonies are found in remote or offshore sites that are difficult to survey consistently, and penguins often nest in burrows or under dense vegetation, rocks, or sand, leading to frequent underestimation of active nests during visual counts.

Notably, African Penguins are highly vocally active during the breeding season, producing frequent display songs linked to mate attraction, territorial defense, and pair coordination. The adult vocal repertoire includes two principal display songs: the Ecstatic Display Song (EDS), produced predominantly by males, and the Mutual Display Song (MDS), a duet between mates. The EDS is a long (5.04 s ± 4.17 s) and structurally variable song composed of multiple syllables (A, B, and C) that vary in frequency and amplitude across inhalation and exhalation phases (Favaro et al., 2014), whereas the MDS beginning with pulsed noise and ending with a low-pitched harmonic segment (Favaro et al., 2014). The EDS syllabic composition varies markedly both within and between individuals (Favaro et al., 2020) and EDS emission is influenced by diel cycle, wind, and temperature (Hacker et al., 2023). In this context, the combination of high vocal activity during the breeding season and the logistical difficulties of visual surveys makes PAM a non-invasive and operationally feasible alternative for continuous monitoring without disturbing the birds (Boersma et al., 2020; Gibb et al., 2019; Sugai et al., 2019).

Building on this foundation, the present study investigates the relationship between nest density and vocal activity, a goal that requires the reliable and scalable detection of penguin vocalizations. We therefore adopt a sequential methodological approach. First, we develop and validate an automated detector for Ecstatic Display Songs using a CNN trained on a heterogeneous dataset that combines *in situ* soundscape recordings from wild colonies with *ex situ* recordings collected under controlled conditions. This strategy is intended to capture the full acoustic variability of the species’ vocal output and background environments, thereby maximizing detector robustness and transferability across space and time. Second, we use the resulting EDS detections to evaluate whether peaks in vocal activity can serve as a quantitative acoustic proxy for local nest density. We hypothesized that areas with higher penguin densities, and thus more active nests, would exhibit correspondingly higher rates of EDS production. By integrating automated acoustic detection with nest census data through statistical modelling, this study assesses the feasibility of PAM as a scalable, low-disturbance tool for estimating nest density. Establishing such a link would provide a robust, transferable, and non-invasive framework to support local conservation monitoring and inform long-term management of this threatened species. Beyond African Penguins, this approach opens new avenues for monitoring a wide range of seabird and terrestrial species, demonstrating how bioacoustics can provide a scalable, sustainable, and minimally invasive framework to tracking wildlife populations in a rapidly changing world.

## 2. Methods

### 2.1 Study area

The study was conducted at the Stony Point Penguin Colony (Betty’s Bay, South Africa), a site of critical conservation importance for the African Penguin. The first penguins established this colony in 1982 after immigrating from declining offshore island populations, and by 2022 Stony Point supported approximately 1,600 breeding pairs, around 15% of the remaining South African population (Masotla et al., 2023).

All research activities were carried out under permits issued by the South African Department of Forestry, Fisheries and the Environment for scientific investigation and experimentation (Res2023-25; Res2024-38; Res2025-13) and by CapeNature (CN32-87-23209; CN32-87-27311; CN32-87-31489). Data collection protocols were also approved by the Ethical Committee of the University of Turin (approval n°0664878) and by the Ethical Committee of the University of the Western Cape (approval n°AR24/1/1).

### 2.2 Acoustic recordings

We used a combination of *in situ* and *ex situ* recordings to capture both the ecological heterogeneity of soundscape from wild breeding colonies and the controlled variability in individual song structure, including high signal-to-noise vocalizations, under captive conditions. This integrated design enhanced detector robustness and intended to improve transferability across sites and environmental contexts.

#### 2.2.1 *In situ* automated recordings

In 2023, acoustic data were collected at eight fixed sampling points within the colony using eight Song Meter Micro recorders (Wildlife Acoustics Inc.). Data collection spanned 77 consecutive days, from February 16 to May 3. The recorders operated continuously from 18:00 to 08:00, capturing vocal activity in 30-minute recordings (Tab. S1). We set the recording schedule to conserve battery and storage, aligning duty cycles with prior evidence on African Penguin vocal activity and their predominantly diurnal foraging (Eggleton & Siegfried, 1979; Favaro et al., 2021). Each unit, designed for long-term outdoor deployment, featured a waterproof polycarbonate casing and an integrated omnidirectional microphone. To ensure stability and minimize environmental interference, the recorders were mounted on 50 cm wooden stakes, positioning the microphones 30 cm above ground level-aligning with the average height of the target species while reducing wind interference. The recording settings were optimized with a sampling rate of 48 kHz. The omnidirectional microphone has a sensitivity of +12 ± 4 dBFS re 1 Pa @ 1 kHz when used with an 18 dB gain setting.

In 2024 and 2025, the data collection period also extended from February to April, and the recording settings mirrored those of 2023 to ensure data consistency. However, unlike the continuous recording approach used in 2023, the 2024 and 2025 protocols targeted peak vocalization periods, with recorders operating in two daily time windows, from 04:00 to 08:00 in the morning and from 18:00 to 22:00 in the evening, the schedule was adjusted to better align with local sunrise and sunset. Ten Song Meter Micro recorders were deployed over an 82-day period (2024) and 76-day period (2025) (Fig. 1). The number of sampling points increased to 93 (2024) and 103 (2025) to enhance spatial coverage (Tab. S1). A systematic sampling approach (Sutherland et al., 2004) was employed, with recorders relocated weekly to different areas of the colony, ensuring equal coverage across the study area. To mitigate wind noise, custom windshields were affixed to the microphone covers, an essential adaptation given the often-windy conditions along the South African coastline.

**Figure 1.**
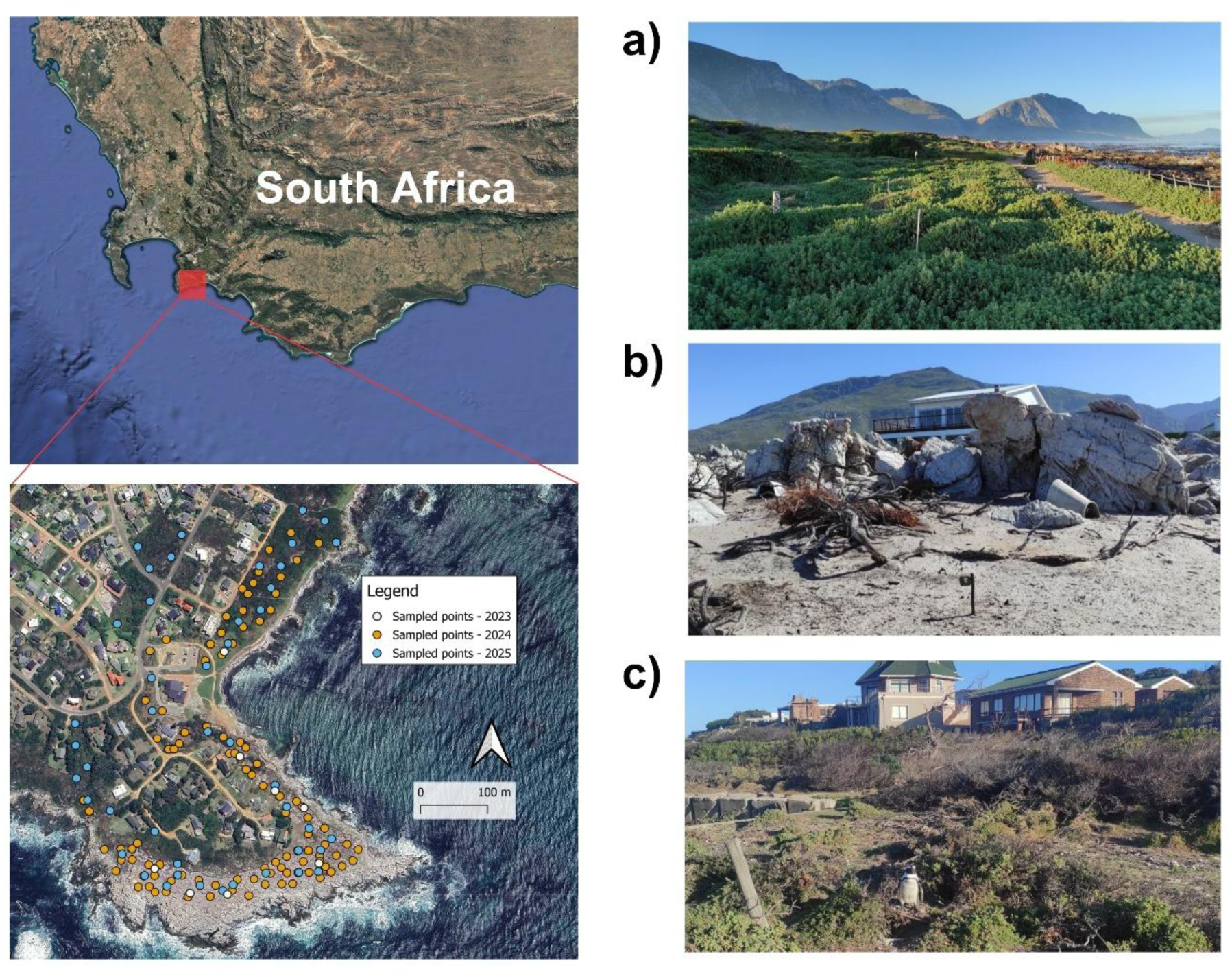
Study area of the Stony Point Penguin Colony (Betty’s Bay, South Africa), showing the locations of acoustic recording points deployed during the 2023–2025 breeding seasons. Right-hand panels illustrate the main habitat types within the colony: (a) areas dominated by dense vegetation; (b) coastal zones characterized by sandy terrain interspersed with rocky substrates; and (c) areas with sparse shrub cover, primarily *Baccharis halimifolia*.

#### 2.2.2 *In situ* focal animal recordings

In addition to the ARUs data, we collected *in situ* recordings using a Sennheiser MKH 416 shotgun microphone (flat frequency response 40 Hz—20 kHz, sensitivity 25 mV/PA1 dB, max SPL 130 dB) connected to a Canon XA40 4K video camera. We employed a focal sampling approach (Altmann, 1974), selecting individual penguins at the Stony Point Penguin Colony that produced EDS at distances between 3 and 10 meters to ensure optimal signal-to-noise ratio. Recordings were captured in uncompressed PCM format at a sampling rate of 44.1 kHz and a resolution of 16 bits. This setup allowed us to capture high-fidelity audio while simultaneously documenting individual identity.

#### 2.2.3 *Ex situ* focal animal recordings

*Ex situ* focal animal recordings collected from the African Penguin colony housed at Zoomarine Italia (Torvaianica, Rome) marine park. Details on the colony and data collection can be found in Calcari *et al*. (2021). Recordings were collected inside the exhibit at distances of 3 to 10 meters from vocalizing individuals, using a RØDE NTG-2 shotgun microphone (frequency response: 20 Hz–20 kHz; maximum SPL: 131 dB) connected to a ZOOM H5 Handy Recorder (48 kHz sampling rate). Audio files were saved in WAV format with 16-bit resolution. The *ex situ* recordings, collected in a controlled zoo environment, offered high-quality vocalizations with minimal background noise.

### 2.3 *In situ* visual counts of penguin nests

Alongside acoustic monitoring, weekly visual counts of active nests were conducted at the Stony Point Penguin Colony under the supervision of CapeNature rangers, in the same sampling points where Song Meter recorders were deployed in 2024 and 2025. In 2023, we deployed eight recorders at predefined locations that remained unchanged throughout the entire breeding season. These recordings were used to confirm the previously reported peak in vocal activity; no nest censuses were conducted during this season.

The primary goal of these surveys was to document nest density of the colony following the standard monitoring protocols used by local rangers. A nest was classified as active when at least one clear indicator confirmed penguin presence, such as fresh guano, recent digging activity, the presence of eggs, or direct observations of adults or chicks inside the nest. Nests showing no evidence of occupation, such as spider webs, old guano, or dense overgrown vegetation, were excluded from the count (Francomano et al., 2024; Fig. S1). In 2024, active nests were counted within square plots of 20 × 20 m (400 m²), following standard CapeNature habitat-assessment protocols. In 2025, nest counts were conducted within circular plots of 10 m radius (314 m²) centered on each recorder. This change was implemented to better align the spatial extent of visual surveys with the effective acoustic footprint of the recorders.

The choice of a 10 m radius was guided by propagation experiments conducted in late 2024 at the study site. These experiments showed that the most distinctive syllables of the EDS, particularly the B syllable, remained reliably detectable up to approximately 12 m under calm conditions. Beyond this distance, vocal identity degraded and acoustic structure weakened, reducing classification confidence (Ninni et al., 2024). Based on these results, a 10 m detection radius was adopted as a conservative estimate of effective acoustic range. This distance defines the spatial domain within which EDS detections and nest counts were assumed to be linked with stable detectability.

### 2.4 Automated detection of African Penguin vocalizations with a Convolutional Neural Network

#### 2.4.1 Data preprocessing and dataset composition

To characterize African Penguin vocal activity, we focused on the EDS. Previous studies have shown that EDS is the most frequently recorded vocalization in passive acoustic recordings, occurring substantially more often than MDS, with an approximate MDS: EDS ratio of 1: 3.4 (Favaro et al., 2021) and primarily produced in a territorial context by one member of a breeding pair while positioned inside the nest. Structurally, the EDS exhibits a stereotyped three-part organization composed of A, B, and C syllables (Fig. 2). The A syllable typically occurs as a series of repeated elements at the onset of the song, with the number of repetitions varying both within and among individuals. The B syllable follows as a single element, or occasionally two, and shows moderate inter-individual variation while remaining relatively stable in its temporal and frequency structure. The C syllable occurs toward the end of the song and exhibits pronounced inter-individual variability and frequency modulation (Favaro et al., 2020). Given this multi-syllabic organization, automated detection was anchored to the B syllable, which provides a consistent and reliable indicator of the presence of a complete EDS while avoiding ambiguity introduced by variable repetition of A syllables and the strong frequency modulation characteristic of C syllables.

**Figure 2.**
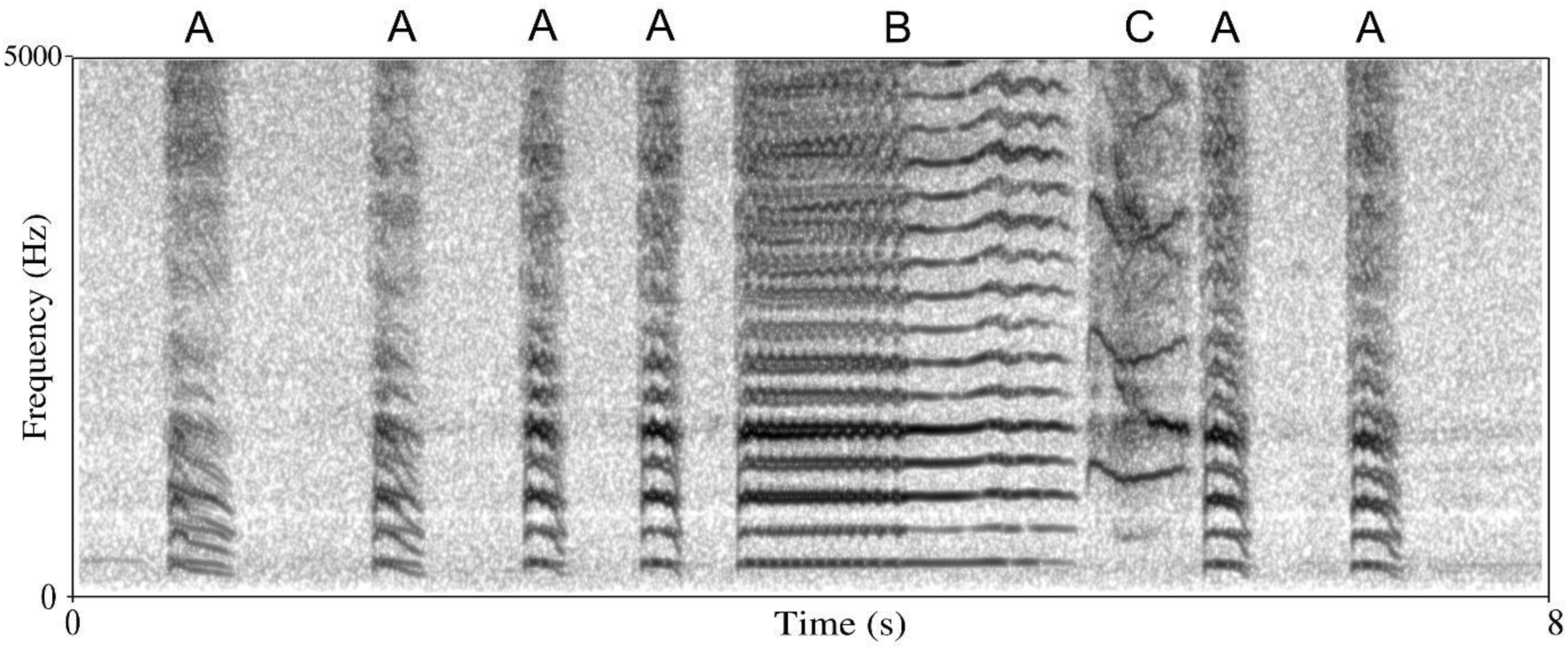
Ecstatic Display Song of an adult African Penguin recorded through Songmeter device at the Stony Point Penguin Colony (A = repeated short syllables; B = long syllable, used to define the positive detection class; C = inspiration syllables). The spectrogram was generated in Praat v. 6.4.34, using a Gaussian window shape, window length = 0.04 s, number of time steps = 1000, number of frequency steps= 500, dynamic range = 70 dB.

To develop the CNN model for detecting EDS, we combined the *ex situ* and the *in situ* recordings. This multi-source approach yielded a diverse acoustic dataset that captured a wide range of environmental conditions, individual variability, and vocalization quality, supporting the development of a robust and scalable model. Data collected *in situ* in 2023 and 2025 were used for model training, whereas those collected in 2024 were reserved for independent testing. This split ensured adherence to the 70–15–15 class-1 ratio. We defined two classes: (i) penguin vocalizations (Class 1; EDS) and (ii) background and non-focal sounds (Class 0; anthropophonies, geophonies, and non-target biophonies. The final EDS dataset comprised 2,453 manually annotated B syllables and 9,678 soundscape segments, each standardized to 1.5 s.

Specifically, soundscape data were selected from a subset (n= 55) of previously annotated long-duration recordings collected in 2023 at the Stony Point Penguin Colony (Terranova et al., 2024), representing a broad range of colony locations and environmental conditions. Recordings were sampled to reflect the acoustic structure of the colony soundscape and were used to construct balanced training and validation datasets. Clips were selected to achieve target proportions of key non-biological and environmental noise sources, including low-, medium-, and high-wind conditions and wave noise. Soundscape data also included non-target biological sounds, such as African Penguin vocalizations other than EDS (e.g. Mutual Display Songs and begging calls) and vocalizations from other species present at the colony (e.g. invertebrates, Cormorants, Kelp gulls, Oystercatchers, and a variety of Passeriformes). To limit temporal autocorrelation, a minimum time gap (5 s) was enforced between clips extracted from the same source recording.

EDS vocalizations were sourced from a combination of *in situ* and *ex situ* recordings. *In situ* EDS were obtained both from passive acoustic monitoring units deployed within the colony and from recordings collected using directional microphones, capturing natural variation in calling behavior. *Ex situ* EDS were drawn from controlled recordings collected outside the colony. Across these sources, the dataset comprised EDS from a total of 135 individual African Penguins, ensuring broad representation of individual-level variation in vocal structure. All EDS were manually annotated at the syllable level, and for the positive detection class, only segments corresponding to the B syllable were retained for model training.

All audio files were downsampled to 16 kHz and converted to fixed-length mono audio segments of 1.5. Segments shorter than their target length were padded with low-level Gaussian noise (σ = 10⁻³), rather than zeros, to avoid hard silence edges and reduce artefacts the model could memorize. We applied time-shifting augmentation with a probability of 50%, whereby the original audio signal was randomly shifted forward or backward in time by up to ±20% of its duration. The shift was implemented by repositioning the signal within the fixed-length input window, while the padding segments were kept unchanged, ensuring that no new acoustic content was introduced. We computed Short-Time Fourier Transform (STFT) spectrograms using a 400-sample window and 160-sample hop, took magnitudes, and applied a log transform to stabilize variance and compress dynamic range. This log-scaling attenuates high-energy background bursts (e.g., wind, waves) and enhances lower-energy, biologically informative structure in penguin vocalizations (Stowell, 2022). Spectrograms were then normalized and expanded to a single-channel tensor for CNN input.

#### 2.4.2 Model architecture, training, and evaluation

The model comprised four sequential convolutional blocks, each including a 2D convolution, batch normalization (Ioffe & Szegedy, 2015), ReLU activation, and a 2×2 max-pooling layer (Su et al., 2019). A SpatialDropout2D layer (0.2) was added after the first block to regularize feature maps and improve robustness to local noise. This four-block configuration provided a balanced trade-off between receptive-field depth and computational efficiency, allowing the model to capture both fine-scale and higher-level spectro-temporal features while limiting overfitting (Abdoli et al., 2019; Su et al., 2019). After convolutional processing, extracted features were condensed through a Global Average Pooling (GAP) layer, followed by two fully connected layers (128 and 64 units) with ReLU activation, L2 regularization, and a final sigmoid neuron outputting the probability of EDS presence.

A structured grid search was performed to explore combinations of learning rates (1×10⁻⁵ – 3×10⁻⁴), L2 regularization strengths (0.001, 0.0005), and dropout rates (20–50%) in the dense layers, using the Adam optimizer and binary cross-entropy loss (Ghani et al., 2023; Kingma & Ba, 2014). Each configuration was trained for up to 50 epochs with early stopping (patience = 10) and adaptive learning-rate reduction (factor 0.5, minimum 1×10⁻⁶).

Model selection followed a hierarchical criterion prioritizing the highest validation ROC-AUC (Receiver Operating Characteristic – Area Under the Curve), followed by the lowest validation loss, highest recall, highest precision, and highest PR-AUC (Precision-Recall). Performance was monitored on the validation set using recall, precision, ROC-AUC, and PR-AUC to assess both discriminative ability and precision–recall trade-offs under class imbalance.

Across the evaluated configurations, the selected model used a learning rate of 3×10⁻⁴, L2 = 0.001, dense dropout = 0.3, and SpatialDropout2D = 0.2 in the first convolutional block. The selected model was then evaluated on an independent test dataset, and the final decision threshold (0.30–0.70 range) was set to the value maximizing the F₁ score.

#### 2.4.3 Quantification and validation of EDS detections in long-duration recordings

The automated model, originally trained on 1.5-second audio windows, was adapted to quantify the number of EDS in continuous recordings. To achieve this, we developed a custom Python pipeline that processes long-duration recordings by dividing them into consecutive 1.5-s segments, applying the trained model to each slice to estimate the total number of EDS per recording. Because a single EDS can comprise multiple B-syllables, which the CNN may detect as separate segments, detections occurring within ≤ 1 s were merged into a single detection event.

As the CNN was trained using high- and medium-quality EDS examples, it is expected to preferentially detect vocalizations produced by individuals close to the recorders, whose calls exhibit higher signal-to-noise ratios and clearer spectral structure. However, soundscape recordings from natural colonies encompass a wide range of vocalization qualities and background noise conditions. We therefore explicitly assessed the performance of automated detection across vocalizations of differing quality levels.

To achieve this, we evaluated the model detections on 316 continuous 30-minute recordings (approximately 150 hours in total) collected in 2024 across multiple recorders and times of day, encompassing a wide variety of acoustic conditions. Each recording was manually annotated by a trained observer, who labelled every EDS and B-syllable and assigned a quality tier based on visual clarity and signal integrity. Vocalizations were classified into three levels: (1) low quality, when signals were faint, noisy, or only partially visible; (2) medium quality, when signals were clear but showed limited harmonic structure or mild interference; and (3) high quality, when signals were prominent, noise-free, and displayed multiple, well-defined harmonics (Fig. S2; Genes et al., 2023; Gridley et al., 2015; Upham-Mills et al., 2020).

We first compared total vocalizations counts from automated detection with manual annotations under three inclusion schemes: (i) high-quality only, (ii) high + medium quality, and (iii) all quality combined. For each annotation tier, we assessed agreement between automated detection and manual counts by computing the Pearson correlation coefficient (r), mean absolute error (MAE), and mean bias (automated counts – manual counts). These analyses evaluated the overall agreement between automated detections and manually annotated EDS across recordings and acoustic conditions. Because these comparisons were based on raw automated detections, which include false positives from automated detection, they capture total detection output rather than true event-level accuracy. The latter was assessed separately through temporal matching of singular detections.

#### 2.4.4 Event-level evaluation of automated detections

We performed an event-based evaluation using temporal matching tailored to the structure of EDS songs (Mesaros et al., 2021). An automated detection was counted as a *true positive* (TP) if it overlapped with a manual annotation within ±0.25 s. Since the model operated at the B-syllable level, a detected B-syllable was considered a TP when its onset and offset occurred entirely within a manually annotated EDS window. Detections not meeting this criterion were classified as false positives (FP), while annotated EDS events without corresponding detections were false negatives (FN). Precision, recall, and F₁-scores were computed using micro-averaging, which first aggregates all detections and annotations across recordings and then calculates the metrics on these global totals (Knight et al., 2017; Mesaros et al., 2021).

### 2.5 Vocal activity and nest density

Once the total number of EDS per 30-minute recording was quantified, we assessed whether vocal activity could serve as a reliable proxy for African Penguin nest density. To test the relationship between vocal activity and nest density, we combined acoustic data and visual nest count data collected during the 2024–2025 breeding seasons, yielding a total of 196 sampled points. For the present analysis, 144 sampling points were retained, each comprising approximately seven consecutive days (2024: 7.9 ± 1.3; 2025: 7.5 ± 1.0) of 30-minute passive acoustic recordings collected at fixed times, alongside concurrent nest counts obtained through field surveys. A small subset of sampling points from the 2025 season was not retained because recordings were obtained using a different acquisition protocol, ensuring methodological consistency across years. We excluded recordings in which wind noise exceeded 70% of the total detection time following the methodology described by Terranova et al. (2024), resulting in the removal of approximately 5% of the acoustic dataset overall (2.67% in 2024 and 2.24% in 2025). This filtering step minimized the impact of high-wind conditions—known to degrade acoustic transmission and classification performance—and helped standardize detectability across sites. Environmental covariates including temperature (°C), relative humidity (%), and wind speed (km/h) were extracted from the nearest weather station (https://www.wunderground.com/hourly/za/betty's-bay), and metadata such as recorder number, time of day, date, and habitat type were annotated for all recordings.

To quantify vocal activity, a commonly used vocal activity index is the Vocal Activity Rate (VAR; Oppel et al., 2014). However, VAR can vary substantially due to short-term behavioral and environmental fluctuations, which limits its reliability as a proxy for abundance. To overcome these limitations, we employed the Maximum Count per Minute (MAX) index (Hutschenreiter et al., 2024), adapted to the longer duration of EDS, hereafter EDS peak. Specifically, for each sampling point, we defined the EDS peak as the highest number of EDS detected among all 30-minute recordings collected at that point. This metric captures the peak intensity of collective calling, a behavioral manifestation of synchronous social activity. By focusing on these periods of heightened vocal behavior, typically associated with mate attraction, territorial defense, or group coordination (Leclerc et al., 2025), the EDS peak provides a more stable and ecologically meaningful vocal activity index in socially vocal species such as the African Penguin.

#### 2.5.1 Statistical Modelling

We fitted Generalized Additive Models (GAMs) using the mgcv package in R (Wood, 2017) to estimate expected nest density as a function of acoustic and environmental covariates. The response variable was the nest density (number of active nests per sampling point), modelled with a Negative Binomial distribution and a log-link function to account for overdispersion. The log-transformed sampled area (m²) was included as an offset to account for differences in the area surveyed among plots. As a robustness check, we refitted the model including plot geometry (square vs circular) as an additional fixed effect, the estimated relationship between EDS peak and nest density remained unchanged. Moreover, because sampling points were spatially structured within the colony, we assessed residual spatial autocorrelation using Moran’s I and evaluated the sensitivity of model inference to spatial correction by refitting a GAM including a two-dimensional smooth of sampling location (Dormann et al. 2007). Because our primary objective was inference on the EDS-nest relationship rather than maximizing spatial prediction accuracy, spatial models were used as sensitivity analyses rather than for primary inference.

The full model included a smooth term for EDS peak, as well as smooth terms for environmental covariates (wind speed, temperature, humidity) and a cyclic spline for time of day to capture diel variation. Recorder identity and calendar date were included as random-effect smooths (bs = “re”) to account for repeated measurements and shared day-to-day variability. Models were fitted using REML.

Collinearity among predictors was assessed using pairwise Pearson correlations, which indicated weak to moderate linear associations (r ≤ 0.36). Because generalized additive models can exhibit nonlinear dependencies not captured by linear correlations, concurvity among smooth terms was also evaluated using the concurvity() function in mgcv (Wood, 2017). High concurvity was observed among temporally structured covariates, reflecting shared seasonal and day-to-day variability that was intentionally absorbed by a calendar date random effect; these terms were therefore retained as controls and not interpreted individually. Model diagnostics were inspected using standard residual plots. These indicated no major deviations from model assumptions and confirmed that the Negative Binomial distribution provided an appropriate fit. Model performance was evaluated against a reduced (null) model excluding the EDS peak term using a likelihood ratio test (χ² test on deviance differences). To identify where the relationship between EDS detections and nest density stabilized, we computed the first derivative of the smooth term for EDS using the derivatives() function in gratia. The plateau was defined as the range of EDS values for which the 95% confidence interval of the derivative included zero, indicating no further significant change in slope.

The final fitted model was used to predict the expected number of active nests across increasing levels of EDS detections. Predictions were generated as a continuous function of EDS while holding all other covariates at reference or mean values. Random effect smooths for date and recorder were excluded during prediction. Predicted values were then summarized within discrete EDS bins to facilitate ecological interpretation, and bin-level medians and 95% model-based confidence intervals were derived from simulations of the fitted model’s coefficient covariance matrix.

To test temporal generalization, a model fitted only on the 2024 data was used to predict nest counts in the independent 2025 data. Predictive performance was evaluated using the coefficient of determination, Spearman’s rank correlation (ρ), root mean squared error (RMSE) and

Mean Absolute Error (MAE), computed between observed and predicted nest counts. Predicted relationships were then visualized, and acoustic activity bins were constructed to summarize observed and predicted nest counts in 2025, with 95% confidence intervals for bin-level means obtained via bootstrap resampling. All analyses were conducted in R version 4.5.0 using the packages mgcv (Wood, 2017), gratia (Simpson, 2024), and tidyverse (Wickham et al., 2019).

## 3. Results

### 3.1 Automated detection of EDS

The deep learning–based detection model for EDS demonstrated high performance in distinguishing penguin vocalizations from complex colony soundscapes. The model achieved a recall of 0.99 and a precision of 0.89, indicating that 89% of positive predictions corresponded to true EDS windows. The AUC reached 0.99, confirming near-perfect discrimination between EDS and background sounds. Threshold optimization was conducted to fine-tune the balance between recall and specificity. The optimal threshold of 0.7 maximized the F_1_-score (0.97). ROC and PR curves further validated these findings, with both curves showing near-perfect discrimination and consistently high precision across all recall levels (Fig. 3 a-b). The confusion matrix reflected this performance, with the model correctly classifying 6,738 background and non-focal sound samples and 419 EDS vocalizations, while misclassifying only 20 false positives and 5 false negatives (Fig. 3c). Overall, these results confirm that the EDS automated detection model achieves highly accurate, robust, and generalizable detection of EDS under variable environmental conditions.

**Figure 3.**
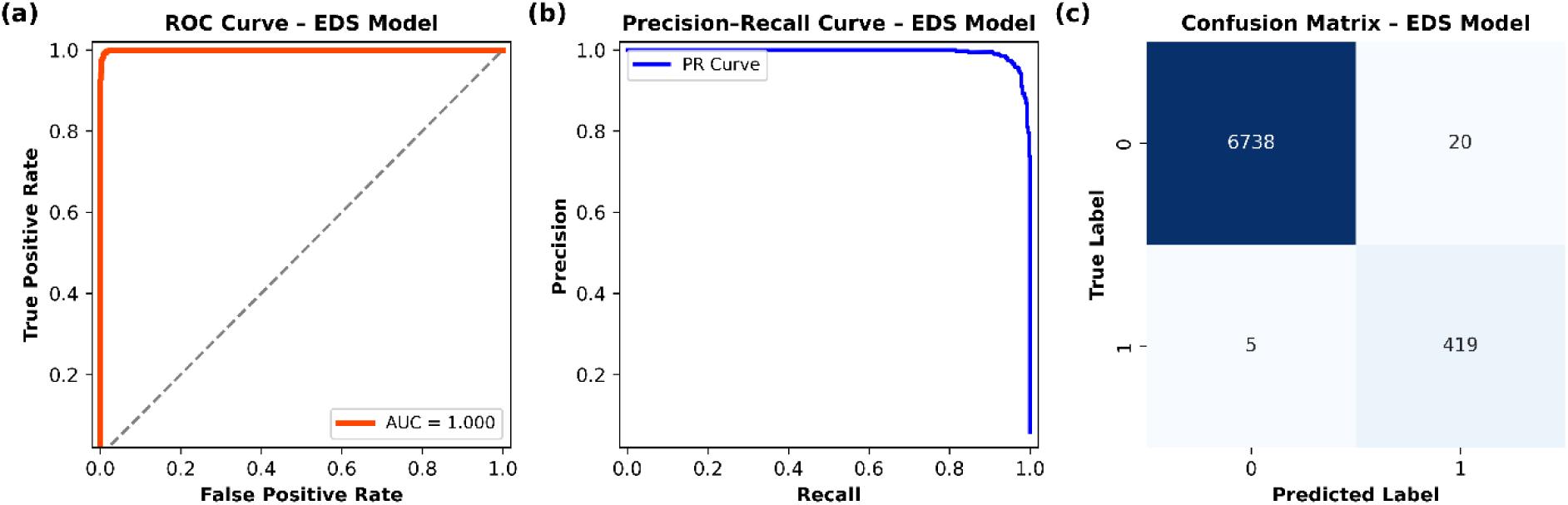
ROC curve (a), PR curve for the automated detection model (b), and Confusion Matrix on the test set (c). The ROC curve (a) shows the trade-off between true positive and false positive rate across thresholds, with an AUC near 1.0 indicating strong classification performance. The PR curve (b) highlights the balance between precision and recall, useful for imbalanced datasets, where higher values suggest better detection of true positives. Class 1 of the Confusion Matrix (c) indicates the target vocalization (EDS) while class 0 indicates non-target vocalizations.

### 3.2 Automated detection performance on long-duration recordings by vocalization quality

Automated detections in long-duration recordings closely matched manual EDS counts across annotation tiers, with correlation coefficients ranging from r = 0.78 to 0.88 (Tab. 1). Agreement was highest when high- and medium-quality annotations were combined (MAE = 4.5; Bias = –1.8).

**Table 1.**
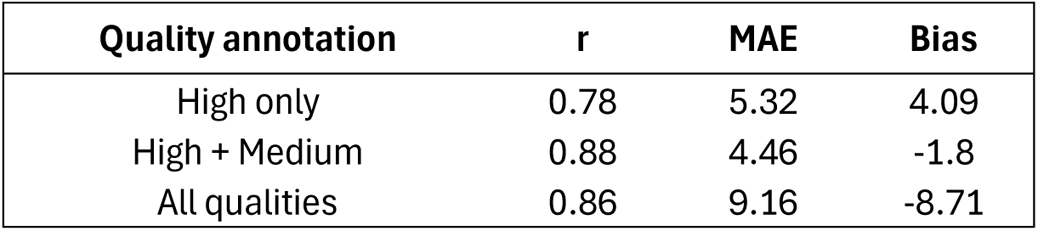
Agreement between automated detection and manual annotations across three annotation-quality levels. The Pearson correlation coefficient (r) quantifies linear agreement, MAE (mean absolute error) measures the average magnitude of absolute deviations between automated and manual counts, and Bias indicates systematic over- or under-estimation (automated − manual). Positive bias values denote automated overestimation while negative values denote underestimation.

| Quality annotation | $r$ | MAE | Bias |
| --- | --- | --- | --- |
| High only | 0.78 | 5.32 | 4.09 |
| High + Medium | 0.88 | 4.46 | -1.8 |
| All qualities | 0.86 | 9.16 | -8.71 |

When comparing the model to high-quality annotations only, automated detections slightly overestimated manual counts (Fig. 4a), likely because the model also captured genuine but fainter EDS that fell below the high-quality threshold used by annotators. In contrast, including low-quality annotations introduced greater uncertainty: the model tended to underestimate manual counts because many low-quality EDS were not detected (Fig. 4c), an expected outcome given that the CNN was trained exclusively on high- and medium-quality vocalizations.

**Figure 4.**
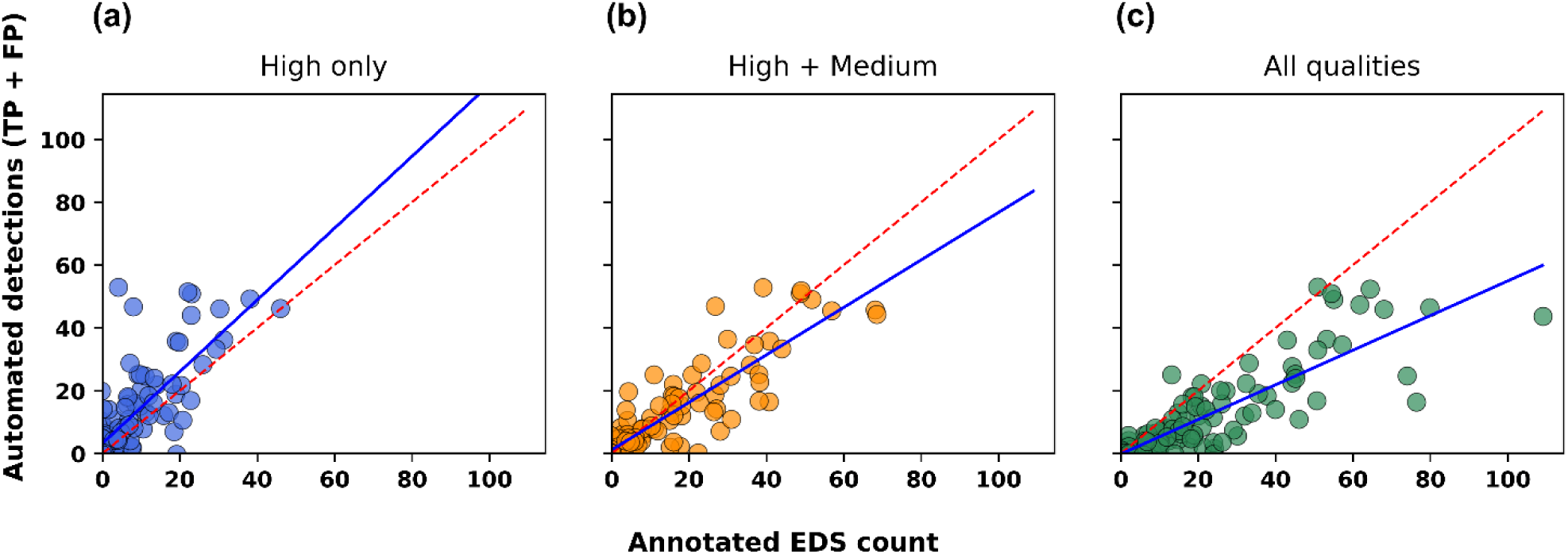
Agreement between manual annotations and automated detections across different EDS quality levels. Each point represents a 30-min recording (n = 108). The red dashed line indicates the 1:1 reference (perfect agreement), and the blue line shows the fitted relationship obtained using ordinary least squares regression (y = ax + b) between total automated detections (TP + FP) and manual annotated EDS counts. Panels correspond to (a) high-quality, (b) high + medium-quality, and (c) all annotation quality levels.

The regressions between automated detections and manual annotations describe how closely total CNN-derived counts track manually annotated EDS counts across recordings. However, even a perfect correspondence in total counts does not guarantee that the model correctly identified the same vocal events. For this reason, we also computed event-based micro-averaged metrics to assess the accuracy of individual detection matches. Among the three annotation schemes, the inclusion of both high- and medium-quality vocalizations yielded the most stable and unbiased performance (Table 2), consistent with the pattern observed in the comparison of raw automated detections and manual annotations. The moderate recall obtained under this inclusion criterion (0.64 on average) likely reflects variability in vocalizations clarity, whereas the higher recall achieved for high-quality annotations highlights the strong influence of signal-to-noise ratio on model sensitivity. To further assess whether apparent false positives reflected genuine but faint EDS rather than spurious detections, we additionally inspected model outputs overlapping low-quality human annotations and reclassified these cases as true positives. Under this corrected scheme, precision reached an overall value of 0.88, indicating that the majority of detections corresponded to real EDS events that were under-annotated due to low signal strength rather than model misclassification.

**Table 2.** Event-based performance of the automated model for detecting EDS under different annotation quality inclusion schemes. Metrics were computed using micro-averaging across all recordings. *Corrected* values account for detections of overlapping low-quality annotated vocalizations, which were reclassified as true positives to avoid penalizing genuine but faint EDS detected.

| EDS annotation quality | Precision | Precision (corrected) | Recall | F1 | F1 (corrected) |
| --- | --- | --- | --- | --- | --- |
| High | 0.46 | 0.88 | 0.76 | 0.57 | 0.82 |
| High + Medium | 0.75 | 0.88 | 0.64 | 0.69 | 0.74 |
| High + Medium + Low | 0.88 | - | 0.48 | 0.62 | - |

Taken together, these validation results indicate that the automated model reliably captures high and medium quality EDS. The strong correspondence with manual counts, supports the use of automated detections to quantify penguin vocal activity within the spatial range at which individuality cues remain preserved. Given known sound propagation limits from playback experiments (∼10–12 m) and the robustness of B-syllables to degradation (Ninni et al.,2024), automated detections can be interpreted as representing callers within a local acoustic neighborhood around each recorder. The validated automated model was therefore applied to the full acoustic dataset for the automated detection of EDS.

### 3.3 Acoustic activity and nest density

Spatial patterns of nest density and vocal activity were highly consistent across breeding areas. As shown in Fig. 5, zones with higher vocal peaks (Fig. 5a) closely overlapped with regions of highest nest density (Fig. 5b).

**Figure 5.**
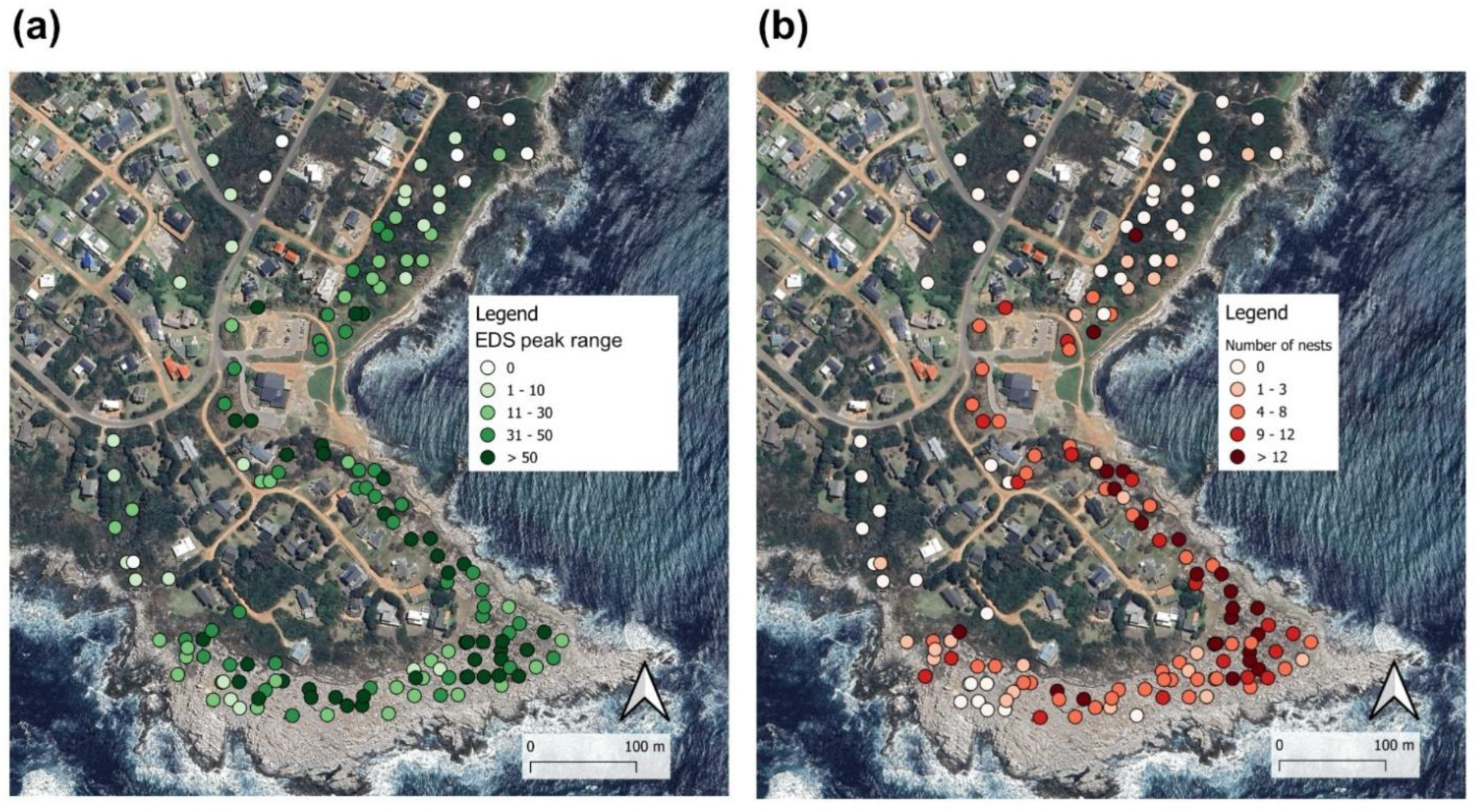
African Penguin vocal peak activity (a) and nest density (b) across the sampling points of the Stony Point Colony during the 2024 and 2025 breeding seasons. Image generated with QGIS software (v. 3.34.4-Prizren).

The modelling analysis reinforced this pattern, revealing that acoustic activity, quantified as the EDS peak, is a strong and nonlinear predictor of nest density. The GAM fitted to the combined 2024–2025 dataset (Negative Binomial family) explained 73.5% of the deviance (adjusted R² = 0.57). The smooth effect of EDS peak was highly significant (edf = 3.42, p < 0.001), whereas meteorological covariates (wind speed, temperature, humidity) and time of day showed no detectable effects (all p > 0.25; Table 3). Including EDS peak significantly improved model fit relative to a reduced model (ΔDeviance = 54.39, Δdf = 1.30, p < 0.001).

**Table 3.** Summary of the GAM. The model includes smooth terms for environmental covariates and random effect smooths for date and recorder, with habitat as a parametric factor. For smoothed terms, edf indicates the complexity of the smooth, Ref.df the reference degrees of freedom, Chi.sq the approximate test statistic, and p-value the significance of the smooth. For parametric coefficients, Estimate is the effect size (on the log scale), Std. Error its uncertainty, z value the test statistic, and p-value the associated significance level.

| Smoothed terms | edf | Ref.df | Chi.sq | p-value |
| --- | --- | --- | --- | --- |
| s(EDS peak) | 3.43 | 3.78 | 67.32 | <0.001*** |
| s(wind_speed_kmh) | 1 | 1 | 0.01 | 0.9 |
| s(Temperature_C) | 2.62 | 2.96 | 4.53 | 0.18 |
| s(Humidity) | 1 | 1 | 0.4 | 0.52 |
| s(time) | 7.5 | 3 | 0 | 0.87 |
| s(date, bs = "re") | 2.6 | 68 | 50.66 | <0.001*** |
| s(recorder, bs = "re") | 4.3 | 9 | 0 | 0.81 |
| Parametric coefficients | Estimate | Std. Error | z value | p-value |
| (Intercept) | -5.24 | 0.18 | -27.03 | <0.001*** |
| Rocks and sand | 0.65 | 0.23 | 2.85 | 0.004** |
| Shrubs | 1.08 | 0.22 | 4.95 | <0.001*** |

Predicted relationships showed a steep increase in nest density at low acoustic activity (< 20 EDS detections per 30 min), followed by a progressive flattening of the curve beyond approximately 44 EDS, indicating saturation of the acoustic index. Derivative analysis of the EDS smooth confirmed this pattern, with the 95% confidence interval of the first derivative overlapping zero from ∼44 EDS onwards, identifying a plateau beyond which additional vocal activity was not associated with further increases in nest density (Fig. 6a). In this saturated range, predicted nest density stabilized at approximately 10–11 nests per sampling site. The observed asymptote likely reflects a detection ceiling: the long duration of EDS calls (5.04 ± 4.17 s), combined with frequent call overlap, constrains the number of distinct detections within a 30-min recording, producing an apparent saturation of the acoustic index in densely populated areas.

**Figure 6.**
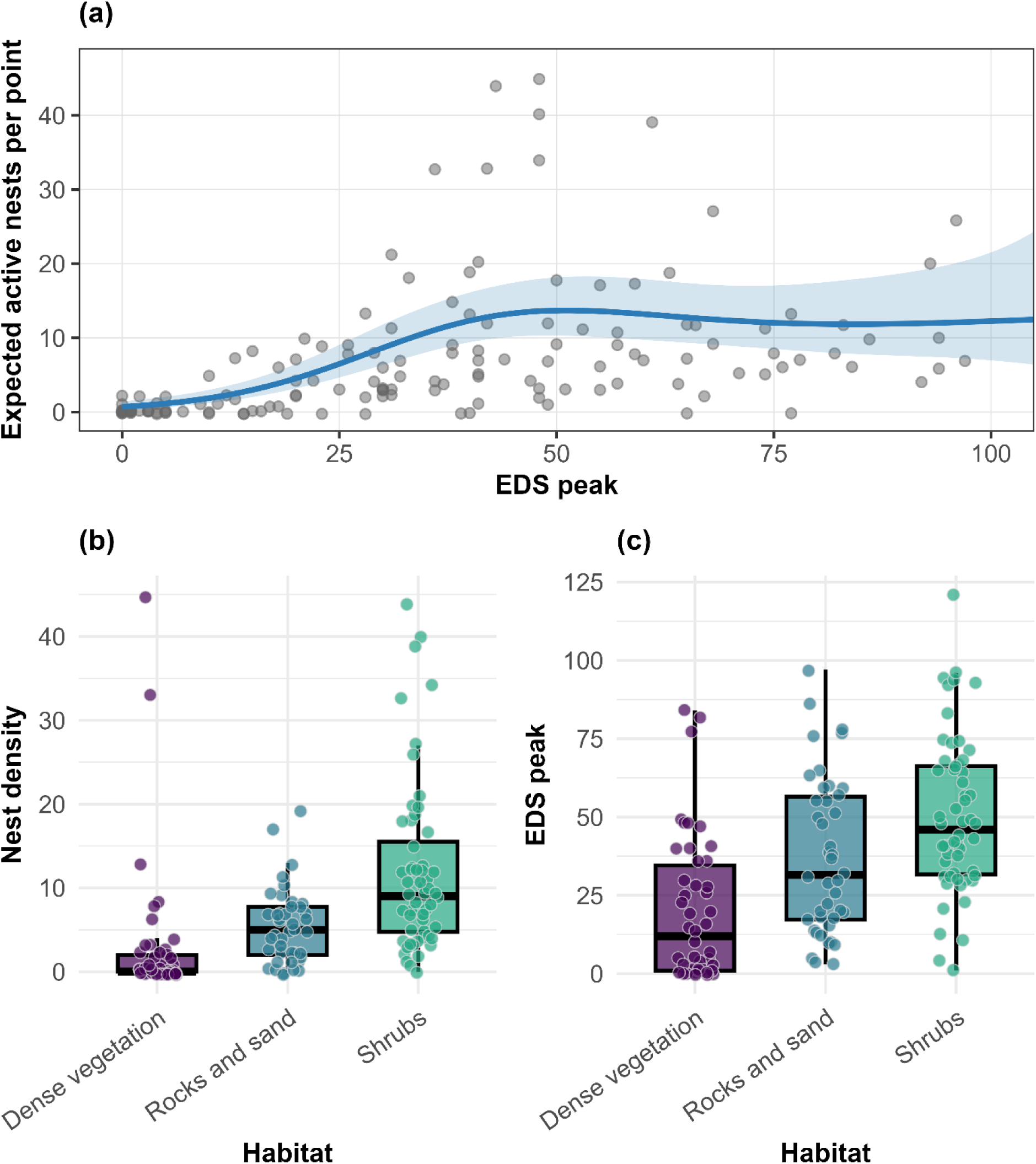
**(a)** Expected number of active nests per 10 m radius as a function of EDS peak detections per 30-min recording, estimated using a Negative Binomial GAM. The line shows fitted values and the shaded band shows the model-based 95% confidence interval; points show observed counts (x-axis displayed from 0–100 EDS for clarity). **(b)** Boxplots showing the distribution of observed active nest counts across habitat types. **(c)** Boxplots showing the distribution of observed EDS peak detections across habitat types.

Calendar date, included as a random intercept, captured substantial day-to-day variability among recordings without altering the estimated EDS–nest relationship. Vocal activity varied among habitats, being lowest in dense vegetation and highest in shrub-dominated areas. This pattern is consistent with the spatial organization of the colony, where nesting areas are clustered within the rocky and shrubby core of the reserve. Residual diagnostics indicated significant spatial autocorrelation in the non-spatial model (Moran’s I, p < 0.01; Table S2), consistent with this clustered configuration. Importantly, when we refitted the model including a two-dimensional smooth of spatial coordinates, residual spatial dependence was removed and the EDS-nest relationship (including its plateau) remained unchanged (Table S3). Accordingly, we report results from the non-spatial model for clarity and parsimony.

Model-based predictions closely aligned with observed nest counts across all EDS peak ranges (Table 4), supporting the use of EDS activity as a robust acoustic proxy for nest density. For interpretability, EDS values were grouped into five activity categories, each corresponding to a distinct range of nest densities, spanning from acoustically inactive areas to saturation at high vocal activity levels.

**Table 4.**
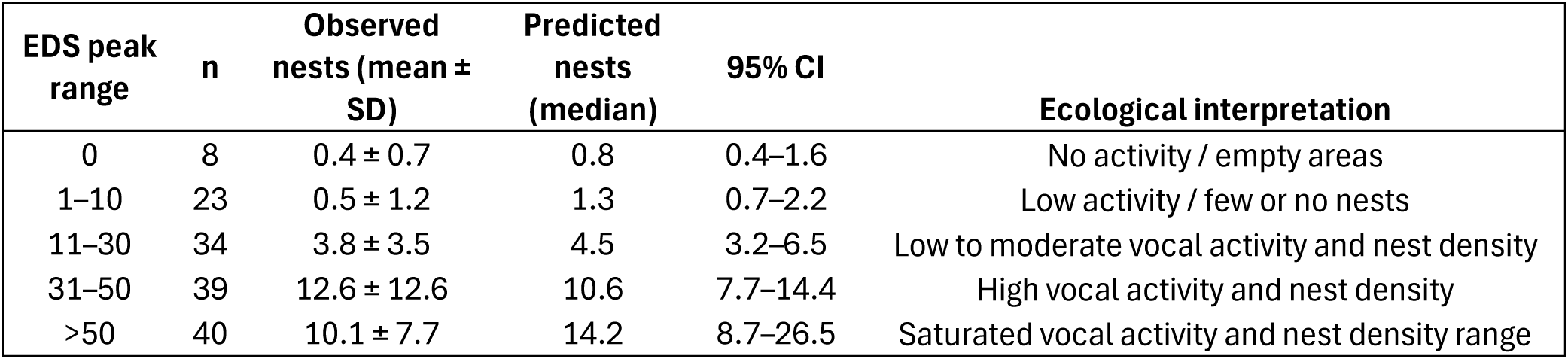
Comparison between observed and model-based estimates of African Penguin nest density across bins of Ecstatic Display Song peak activity. For each EDS peak range, the table reports the number of sampling points (n), the mean ± SD of visually observed active nests, and the median predicted nest density with 95% model-based confidence intervals derived from the GAM. The ecological interpretation summarizes the expected colony state associated with increasing levels of vocal activity, highlighting the nonlinear increase and saturation of the vocal–nest relationship at high EDS levels.

The model trained on 2024 data generalized well to the independent 2025 dataset (R² = 0.50, ρ = 0.77, RMSE = 4.52 nests, MAE = 2.78 nests), demonstrating strong temporal consistency between EDS peak and nest density (Table S4). The total predicted number of nests (232) was within 10.8 % of the observed colony total (209). Mean observed and predicted nest counts were broadly consistent across EDS bins in 2025, with overlapping 95% bootstrap confidence intervals in most bins (Fig. 7a). Together with the observed–predicted agreement at the plot level (Fig. 7b), this supports the temporal transferability of the model trained on 2024 data to the independent 2025 dataset.

**Figure 7.**
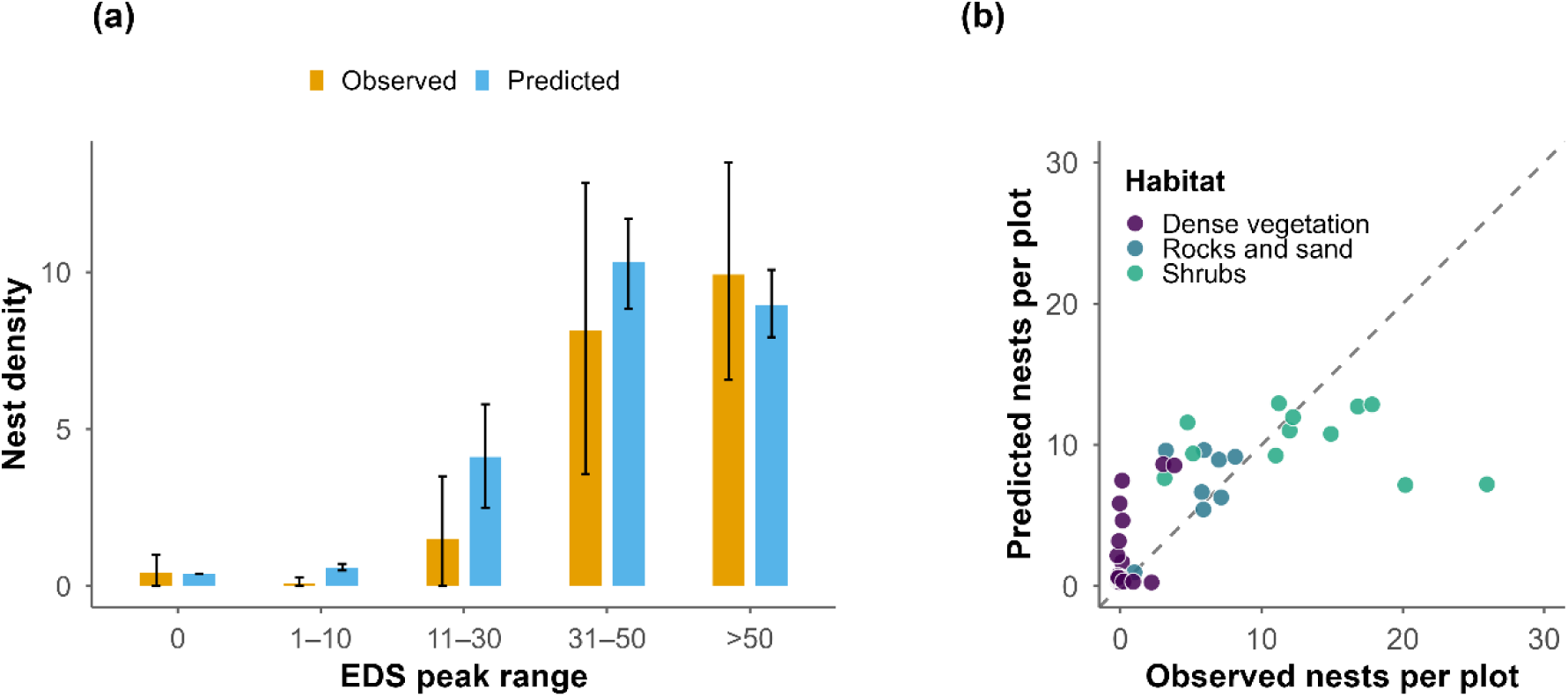
Year-to-year robustness of the EDS–nest density model. Model trained of 2024 data and tested on 2025 data. **(a)** Observed and predicted mean nest density across EDS peak bins for the 2025 data. Error bars show 95% bootstrap confidence intervals. **(b)** Predicted versus observed nest counts for the 2025 dataset (n = 99) using the model trained on 2024 data. Points are colored by habitat, and the dashed line represents the 1:1 relationship.

## 4. Discussion

This study implemented an integrated acoustic monitoring framework that combines large-scale deployment of autonomous recording units with a deep-learning detector for Ecstatic Display Songs to estimate nest presence and density in a wild African Penguin colony. By linking automated vocal detections to ground-truth nest counts, we show that non-invasive acoustic monitoring can provide colony-wide estimates of nest density that are reproducible across years and scalable to large spatial extents because additional spatial coverage can be achieved by deploying more recorders without increasing observer time or disturbance.

Leveraging pre-segmented vocalization samples from both *ex situ* (zoo) and *in situ* datasets, the custom-built convolutional model achieved high detection accuracy across acoustically diverse environments. The inclusion of high-quality *ex situ* recordings enhanced the representation of inter-individual variation in EDS and improved model generalization, consistent with recent advances in deep-learning bioacoustics that automate detection in complex soundscapes (Dufourq et al., 2022; Ruan et al., 2022; Stowell, 2022; Tuia et al., 2022). This integration substantially reduced the need for manual annotation, streamlining data processing and enabling large-scale ARUs deployment.

From a remote-sensing perspective, these findings underscore the value of acoustic sensors as proximal remote-sensing systems capable of detecting biological activity that cannot be reliably captured by satellite or aerial imagery. Vegetation density, terrain obstruction, and the burrow-nesting behavior typical of penguins often hinder visual nest counts, yet these constraints do not impede acoustic propagation. By integrating ARU-derived acoustic measurements with deep learning, we demonstrate a scalable and automated remote-sensing framework capable of extending monitoring capabilities into visually inaccessible habitats. Importantly, the stability of the EDS Peak–density relationship across multiple breeding seasons suggests that acoustic remote sensing can provide temporally robust indices suitable for long-term population surveillance. In this way, acoustic sensing complements traditional remote-sensing approaches by capturing individual vocal activity that directly informs colony-level ecological patterns.

Our results also align with a growing body of evidence demonstrating that vocal activity indices can serve as reliable, cost-efficient proxies for estimating population size and breeding status across avian taxa (Borker et al., 2014; Francomano et al., 2024; Oppel et al., 2014; Pérez-Granados & Traba, 2021). Specifically, we show that acoustic activity can effectively track the nest density of African Penguins, supporting earlier conclusions that passive acoustic approaches can match or surpass the accuracy of traditional census techniques (Boullhesen et al., 2021; Castro et al., 2019; Darras et al., 2019; Digby et al., 2013; Leach et al., 2016).

Nest presence significantly increased vocal activity, with active sites exhibiting ∼75% more EDS than non-breeding areas. However, the relationship between vocal activity and nest density was non-linear. EDS detections rose steeply with nest number until reaching an asymptote at high densities—around 44 EDS per 30 minutes, corresponding to ∼10–11 nests per site. This plateau likely reflects acoustic saturation, where long EDS duration and overlapping songs limit the number of distinct detections. Similar saturation effects have been reported in other colonial species and align with theoretical expectations that acoustic indices rise rapidly at low densities but plateau as masking and temporal congestion constrain detectability (Marques et al., 2013). In the most populated areas of the colony, rapid vocal turn-taking may produce a stable maximum number of detectable EDS even as caller identity changes, meaning the acoustic index saturates before nest density reaches its maximum. Applying individual-recognition techniques could further clarify how caller turnover contributes to the observed plateau and refine estimates in high-density areas.

The temporal stability of this relationship, confirmed by strong generalization from the 2024 model to the independent 2025 dataset, is grounded in the behavioral ecology of the species: EDS are produced almost exclusively by breeding adults engaged in territory defense and courtship, creating a biologically meaningful “one caller–one nest” correspondence. By anchoring vocal activity indices to nest counts rather than individual identity, our approach provides a temporally stable and minimally invasive indicator of nest density across seasons, extending previous observations linking EDS detections to active callers (Hacker et al., 2023).

Generalized Additive Models were central to quantifying the nonlinear relationship between EDS detections and nest density. Their flexibility allowed us to identify the saturation threshold that linear methods would have missed, and their performance is consistent with the GAM-based approach widely applied in abundance and density modelling across diverse taxa (Christensen et al., 2019; Karunarathna et al., 2024; Drexler & Ainsworth, 2013; Frasier et al., 2021; Kosicki, 2020; Clarke et al., 2003). This approach allowed us to clearly visualize and interpret the shape of the relationship, which is critical for understanding ecological processes. However, because the model is based on observed data patterns, predictions beyond the measured EDS range should be treated with caution. In future studies, Random Forest models could help improve predictive performance across more varied conditions by capturing complex, nonlinear interactions—particularly when expanding to multiple colonies where habitat differences introduce additional variability. While less interpretable, these models may offer greater flexibility and robustness when scaling up across spatially and acoustically heterogeneous environments (Kosicki, 2020).

Spatial analyses revealed substantial acoustic activity even in areas visually classified as ‘no-nest’ zones. This pattern was most pronounced in dense vegetation, where areas with few or no visually detected nests nonetheless exhibited high vocal activity (Fig. 6b–c). Such mismatches likely reflect under-detection in visual surveys, as dense vegetation obscures burrow nests, whereas microphones can still capture vocalizations from nearby breeding individuals. Similar detection biases in visually complex environments have been documented in other vertebrate taxa (Marques et al., 2013; Walsh & White, 1999; Waltert et al., 2020). This mismatch underscores one of PAM’s strongest advantages: its capacity to detect cryptic individuals where direct observation fails, reducing habitat-related detection biases that often limit visual surveys.

Despite the strong performance of the EDS-nest density model, some limitations should be acknowledged. First, EDS detections predominantly reflect the vocal activity of breeding adults, which constitute most of the callers within the colony. While the presence of silent breeders cannot be entirely excluded, such cases are likely rare. In contrast, non-breeding individuals, such as sub-adults or adults not engaged in reproduction, typically vocalize little or not at all and are therefore unlikely to be detected acoustically. Consequently, our estimates quantify the density of acoustically active breeding individuals rather than the total number of birds present. Importantly, this limitation mirrors that of standard visual census methods, which likewise exclude non-breeding individuals and report population size in terms of breeding pairs rather than total individuals (Crawford et al., 1990; Kemper 2007).

Second, detectability can be influenced by environmental noise, particularly wind and wave intensity, which affects the propagation and recording of vocalizations in exposed coastal habitats. To address this, we removed recordings with high wind levels from the dataset. While this reduced wind-related interference, wave noise remained and may have continued to affect sound propagation, introducing short-term fluctuations in the recorded vocal activity that do not reflect actual biological patterns. To reduce this source of noise, each sampling site was monitored for six consecutive days, a duration previously shown to stabilize vocal activity estimates (Pérez-Granados et al., 2019; CV ∼ 30%). In addition, the use of the EDS peak as our vocal activity index further mitigates environmental bias. Penguins tend to vocalize most actively under favorable environmental and social conditions. By focusing on the maximum number of EDS detections per point, our approach likely isolates these optimal calling periods, minimizing the influence of transient environmental effects. This explains why covariates such as wind speed, temperature and humidity, factors known to influence vocal behavior (Hacker et al., 2023), did not emerge as significant predictors in our model. In this sense, the EDS peak provides an ecologically meaningful measure of the colony’s readiness to vocalize under its best acoustic and social conditions, enhancing the robustness of the index across variable field settings. Finally, our analysis was restricted to a single colony over two breeding seasons. While this design provided robust temporal validation, applying the same workflow to additional colonies with contrasting densities, topographies, and vegetation structures will be essential for testing its generality and refining colony-specific detection thresholds.

Beyond methodological innovation, this study demonstrates the conservation potential of AI-assisted PAM for long-term monitoring of African Penguins. The capacity to detect penguins in visually obstructed habitats underlines its value as a complement to established methods such as direct observation and mark–recapture using Passive Integrated Transponders (and previously flipper bands). Traditional monitoring approaches such as distance sampling or direct visual nest counts often underestimate true colony size in social and vocally active species, where detection probability varies with habitat complexity, observer distance, and weather conditions (Marques et al., 2013). In contrast, vocal activity indices integrate signals over extended periods, capturing activity even under low visibility. This capacity to account for both temporal and spatial variation in detectability underscores the added value of PAM for long-term and standardized population assessment. Instead, tagging techniques, from flipper bands to Passive Integrated Transponders (PIT tags), have advanced population tracking but can impose stress or logistical constraints (Carroll et al., 2016; Dann et al., 2014; Le Maho et al., 2011; Saraux et al., 2011; Jackson & Wilson, 2002). PAM offers a non-invasive alternative that can be scaled to entire colonies without disturbing birds or altering behavior. Future research should expand ARUs deployments across multiple colonies and integrate fully automated AI pipelines to increase scalability (Sharma et al., 2023; Sugai et al., 2019).

Given the ongoing decline of the African Penguin population (BirdLife International, 2024; Sherley et al., 2024), such integrative and low-impact monitoring approaches are urgently needed. Quantifying the link between acoustic activity and nest density provides a transferable framework that can directly support local conservation efforts. In particular, the method offers rangers and reserve managers a novel tool to monitor penguin presence and nest density across colonies, including in dense vegetation areas where visual surveys are less reliable and mark–recapture or PIT-tagging approaches can be logistically challenging or intrusive. Transforming passive soundscape data into biologically meaningful indicators of colony presence and nest density enables standardized, repeatable, and minimally invasive monitoring at the colony scale. More broadly, the methodological pipeline developed here can be readily adapted to other penguin colonies and species and extended to a wide range of other colonial seabirds, providing a generalizable model for linking acoustic behavior with population processes across taxa.

## Supporting information

Supplementary_materials_doc

## Acknowledgements

We are grateful to the rangers of the Stony Point Nature Reserve for their assistance during the fieldwork and particularly to Gavin Sean Petersen, Cairestine Lottridge, and Marcelin Barry. We also thank Chiara Calcari, Renske Kerkhofs, Franziska Hacker, Nycolin Ninni, Ilaria Morandi, Ines Leclerc, and Linda Gindri for contributing to the data collection and visual inspection of spectrograms. Finally, we acknowledge Zoomarine Rome and particularly Cristina Pilenga.

## Fundings

F. T. was supported by a PON PhD research scholarship (REACT-EU FSE DM 1061). L.T. and X.F. received support from an Erasmus+ Traineeship grant provided by the University of Turin during the fieldwork period. D.R. was partially supported by the National Geographic Society (grant no. NGS-67401R-20). D.R. and N.M. were also supported by the Centre National de la Recherche Scientifique (CNRS), the University of Saint-Etienne, the Institut universitaire de France, and the Labex CeLyA. L.F. was supported by the EMBRC-UP project n° IR0000035, CUP C63C22000570001 entitled “Unlocking the Potential for health and food from the seas”. Field equipment was acquired via the FundsToGether program of the University of Turin (“*Salviamo il Pinguino Africano*” - grant no. FAVL_RIC_N_COMP_24_01)

## Data and code availability

All data and scripts associated with this manuscript are available on Zenodo (https://doi.org/10.5281/zenodo.17778605).

## Notes

https://doi.org/10.5281/zenodo.17778605

### Competing Interest Statement

The authors have declared no competing interest.

