## Supplementary_materials_doc for "Acoustic remote sensing with deep learning enables non-invasive estimation of seabird nest density"

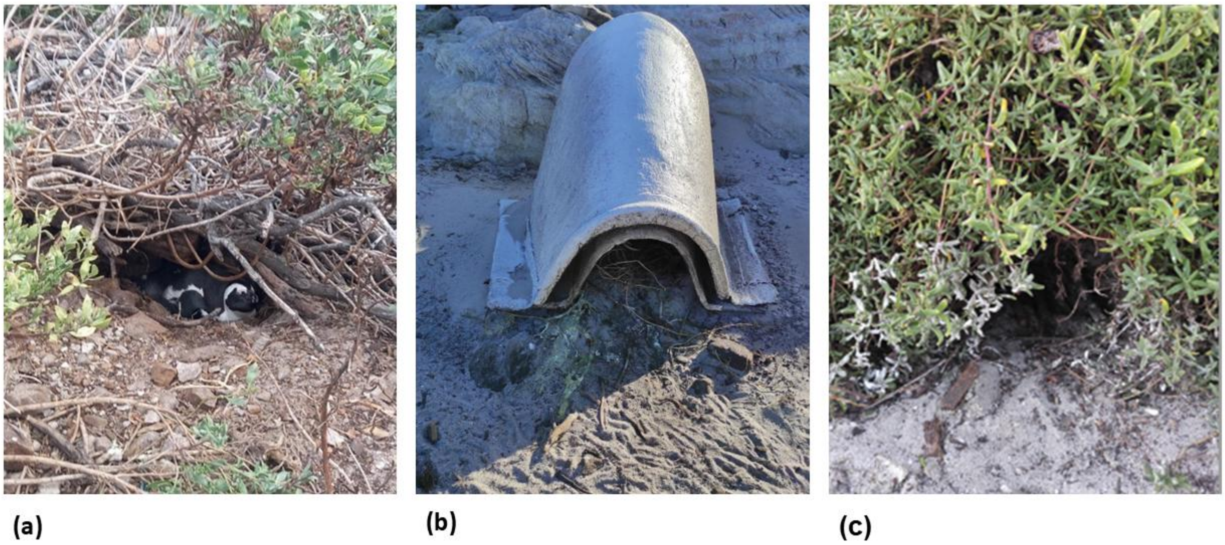


**Figure S1.** Active nests at the Stony Point Penguin Colony (a) showing footprints (b) and guano traces (c).

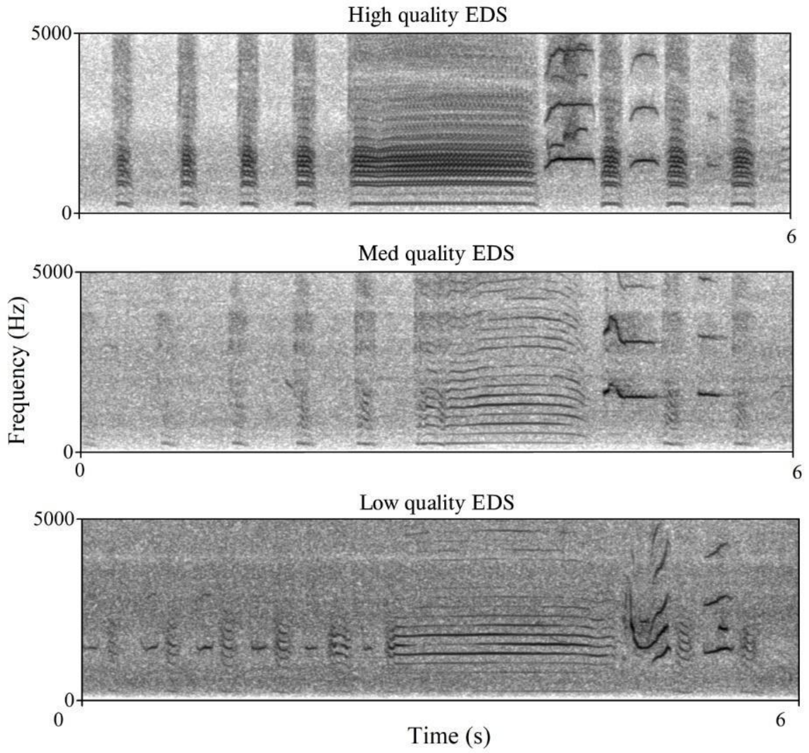


**Figure S2.** Spectrograms of EDS illustrating the three quality categories used in this study (high, medium, and low). Each example shows the temporal and spectral structure characteristic of its respective category, highlighting differences in signal clarity, background noise, and overall detectability. The spectrogram was generated in Praat v. 6.4.34, using a Gaussian window shape, window length = 0.04 s, number of time steps = 1000, number of frequency steps = 250, dynamic range = 70 dB.

**Table S1. Summary of acoustic data collection across the three breeding seasons (2023–2025).** The table reports the total number of recording days, use of a rotating recorder deployment scheme, number of sampling sites, total recordings obtained, and percentage of recordings removed due to wind noise.

| Year | Recording days | Rotation scheme | Sampling sites (n) | Recordings (n) | Windy rec removed (%) |
|---|---|---|---|---|---|
| 2023 | 77 | no | 8 | 12,199 | 15.90% |
| 2024 | 84 | yes | 93 | 13,149 | 2.67% |
| 2025 | 76 | yes | 103 | 8,308 | 2.24% |

**Table S2. Moran's I statistics assessing residual spatial autocorrelation for the non-spatial and spatially adjusted GAMs.** Moran's I was computed on model residuals across a range of neighborhood sizes (k = 5–10). Significant positive spatial autocorrelation was detected in residuals of the non-spatial model, whereas inclusion of a two-dimensional spatial smooth effectively removed residual spatial dependence, yielding non-significant Moran's I values across all neighborhood definitions.

| k | I | expectation | variance | p_value | model |
|---|---|---|---|---|---|
| 5 | 0.122997 | -0.007092199 | 0.00241919 | 0.012 | non_spatial |
| 6 | 0.121317 | -0.007092199 | 0.00198191 | 0.002 | non_spatial |
| 7 | 0.115436 | -0.007092199 | 0.00168842 | 0.005 | non_spatial |
| 8 | 0.118279 | -0.007092199 | 0.00145604 | 0.002 | non_spatial |
| 9 | 0.121548 | -0.007092199 | 0.00127097 | 0.002 | non_spatial |
| 10 | 0.116891 | -0.007092199 | 0.00113174 | 0.002 | non_spatial |
| 5 | -0.03183 | -0.007092199 | 0.00246274 | 0.697 | spatial |
| 6 | -0.02824 | -0.007092199 | 0.00201757 | 0.658 | spatial |
| 7 | -0.02441 | -0.007092199 | 0.00171879 | 0.631 | spatial |
| 8 | -0.01717 | -0.007092199 | 0.00148221 | 0.564 | spatial |
| 9 | -0.01715 | -0.007092199 | 0.0012938 | 0.562 | spatial |
| 10 | -0.01843 | -0.007092199 | 0.00115205 | 0.575 | spatial |

**Table S3. Comparison of key parameters from non-spatial and spatially adjusted GAMs.** Reported metrics summarize the shape, strength, and saturation of the EDS smooth, as well as overall model performance.

| Term | Non-spatial model | Spatial model |
|---|---|---|
| s(EDS peak) edf | 3.42 | 3.28 |
| Plateau onset (EDS) | ~44 | ~43 |
| Deviance explained | 73.50% | 80.70% |
| $\chi^2$ of s(EDS peak) | 54.95 | 31.88 |
| p-value of s(EDS peak) | < 0.001 | < 0.001 |
| Habitat (Shrubs) | significant | significant |

**Table S4. Comparison between model-predicted and observed nest counts across sampling sites differing in habitat type. Predictions were generated for the 2025 data using a model trained on the 2024 dataset.** Each row represents a sampling point at Stony Point Penguin Colony (2025 dataset), showing the habitat category, the EDS peak detected per 30-minute recording, and the corresponding number of nests predicted by the 2024 EDS–nest density model versus those observed during visual surveys.

| ID | Habitat | EDS peak | Predicted nests | Observed nests |
|---|---|---|---|---|
| 619 | Dense vegetation | 77 | 5.89 | 0 |
| 335 | Dense vegetation | 40 | 7.65 | 0 |
| 745 | Dense vegetation | 28 | 4.49 | 0 |
| 660 | Dense vegetation | 23 | 3.19 | 0 |
| 71 | Dense vegetation | 19 | 2.33 | 0 |
| 29 | Dense vegetation | 14 | 1.51 | 0 |
| 187 | Dense vegetation | 7 | 0.79 | 0 |
| 311 | Dense vegetation | 5 | 0.65 | 0 |
| 741 | Dense vegetation | 5 | 0.65 | 0 |
| 161 | Dense vegetation | 3 | 0.53 | 0 |
| 786 | Dense vegetation | 3 | 0.53 | 0 |
| 26 | Dense vegetation | 2 | 0.48 | 0 |
| 24 | Dense vegetation | 1 | 0.44 | 0 |
| 266 | Dense vegetation | 1 | 0.44 | 0 |
| 314 | Dense vegetation | 1 | 0.44 | 0 |
| 870 | Dense vegetation | 1 | 0.44 | 0 |
| 396 | Dense vegetation | 0 | 0.4 | 0 |
| 437 | Dense vegetation | 0 | 0.4 | 0 |
| 868 | Dense vegetation | 0 | 0.4 | 0 |
| 908 | Dense vegetation | 0 | 0.4 | 0 |
| 995 | Dense vegetation | 0 | 0.4 | 0 |
| 1040 | Dense vegetation | 0 | 0.4 | 1 |
| 296 | Rocks and sand | 9 | 1.05 | 1 |
| 74 | Dense vegetation | 0 | 0.4 | 2 |
| 424 | Dense vegetation | 48 | 8.67 | 3 |
| 383 | Rocks and sand | 55 | 9.48 | 3 |
| 208 | Shrubs | 30 | 7.53 | 3 |
| 535 | Dense vegetation | 47 | 8.61 | 4 |
| 982 | Shrubs | 74 | 9.27 | 5 |
| 321 | Shrubs | 41 | 11.69 | 5 |
| 769 | Rocks and sand | 76 | 6.65 | 6 |
| 512 | Rocks and sand | 55 | 9.48 | 6 |
| 425 | Rocks and sand | 30 | 5.6 | 6 |
| 898 | Rocks and sand | 78 | 6.41 | 7 |
| 599 | Rocks and sand | 60 | 8.93 | 7 |
| 641 | Rocks and sand | 59 | 9.06 | 8 |
| 541 | Shrubs | 74 | 9.27 | 11 |
| 324 | Shrubs | 53 | 12.91 | 11 |
| 984 | Shrubs | 66 | 10.83 | 12 |
| 1024 | Shrubs | 42 | 11.95 | 12 |
| 628 | Shrubs | 38 | 10.74 | 15 |
| 849 | Shrubs | 55 | 12.74 | 17 |
| 937 | Shrubs | 50 | 13 | 18 |
| 896 | Shrubs | 93 | 7.24 | 20 |
| 804 | Shrubs | 96 | 7.14 | 26 |
