## Supplementary figures and images for "Acoustic remote sensing with deep learning enables non-invasive estimation of seabird nest density"

### Figure_S1

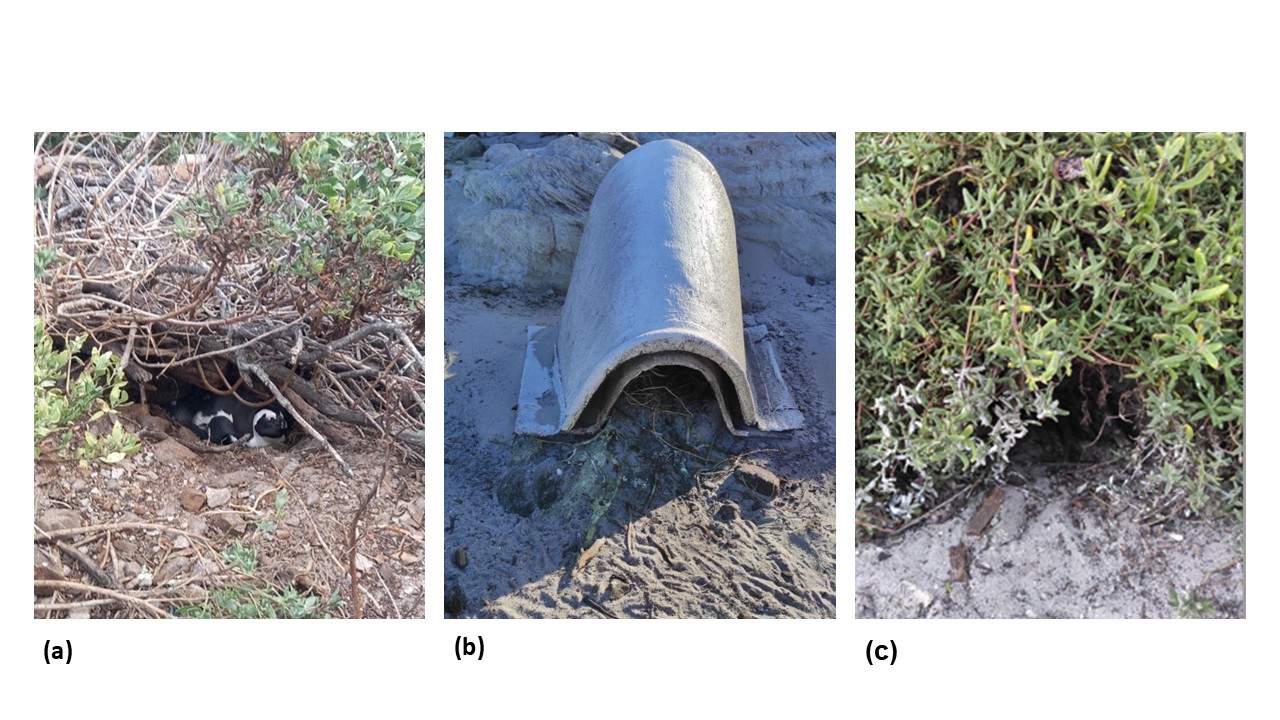

### Figure_S2

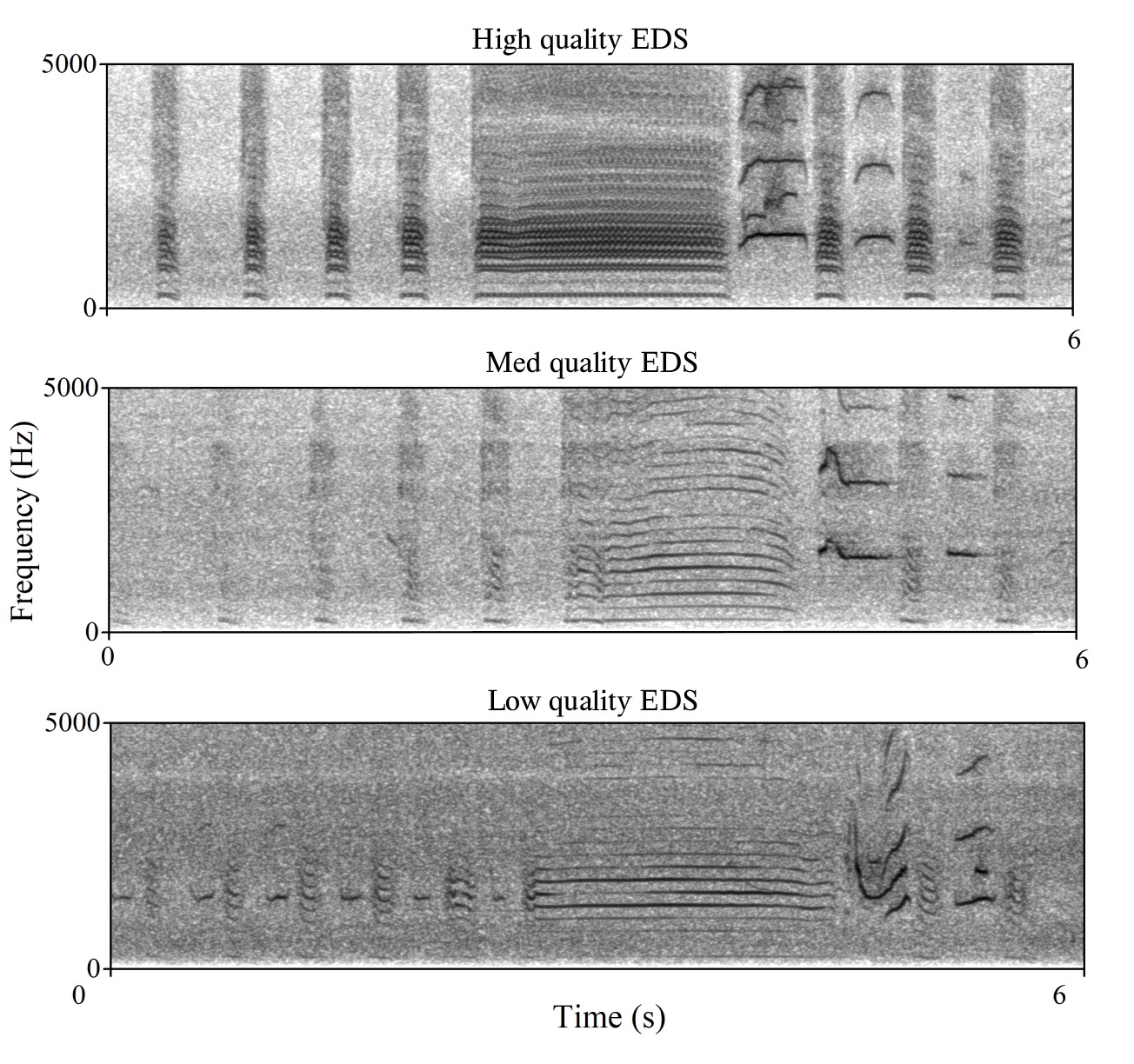
